# Intranuclear Quantum Sensing with Fluorescent Nanodiamond Enabled by Electron-Irradiation and Surface-Chemical Optimization for Microinjection

**DOI:** 10.64898/2026.04.22.720257

**Authors:** Yuki S. Kato, Kota Shiraya, Yukiho Shimazaki, Anna Gatz, Daisuke Fujimaki, Hiroshi Abe, Takeshi Ohshima, Keisuke Fujita, Yoshie Harada, Shingo Sotoma

## Abstract

Fluorescent nanodiamonds (FNDs) containing nitrogen-vacancy (NV) centers are promising quantum sensors for intracellular measurements, yet nuclear applications remained out of reach because optically detected magnetic resonance (ODMR) signals are weak and capillary delivery is inefficient. This study addresses both constraints by optimizing the electron irradiation dose to balance NV creation and charge-state stability, and by grafting hyperbranched polyglycerol with terminal carboxyl groups (HPGCOOH) to suppress aggregation and prevent needle clogging. The optimized dose yields strong ODMR contrast while preserving fluorescence suitable for microscopy. HPGCOOH surfaces enable smooth and reproducible microinjection through fine capillaries. Using this strategy, the microinjection of ODMR-active FNDs into the nuclei of living COS7 cells is achieved, and clear intranuclear spectra comparable to cytoplasmic readouts are obtained. Furthermore, field-of-view temperature sensing across multiple cell nuclei is demonstrated, enabling quantitative and spatially resolved thermal mapping within the genomic environment. This methodology provides a practical route to nuclear quantum sensing and opens opportunities for nanoscale physicochemical measurements within the genomic environment.

## 1. Introduction

Quantum sensing using solid-state spin defects has opened new possibilities for visualizing and quantifying physical and chemical processes in living systems [1]. Fluorescent nanodiamonds (FNDs) embedded with nitrogen-vacancy (NV) centers have attracted attention due to their unique spin-dependent fluorescence properties [2-4]. NV centers enable optically detected magnetic resonance (ODMR) and spin relaxometry, providing precise and spatially resolved readout of physicochemical parameters such as temperature and magnetic fields [5]. Conventional fluorescent probes for intracellular physicochemical sensing based on small molecules or proteins often suffer from photobleaching and signal fluctuations caused by local environmental changes, which can compromise quantitative stability and calibration robustness in live-cell measurements. In contrast, FNDs offer exceptional photostability [2] and physicochemical robustness[6], supporting reliable, long-term and spatially resolved sensing with subcellular resolution [7, 8]. These features make FNDs highly promising for analyzing dynamic and complex processes inside living cells.

FND-based quantum sensing has demonstrated successful intracellular measurements of diverse physicochemical parameters, including temperature[9-14], radical species[15-19], and local rheological properties [20-23]. For example, intracellular thermometry revealed subcellular thermogenesis during neuronal differentiation [14]. In addition, relaxometry enabled minute-scale, real-time tracking of a local redox reaction by sensing time-dependent changes in magnetic noise from polymer-bound nitroxide radicals [15]. As these sensing techniques have advanced, the field has begun to shift toward site-specific and organelle-targeted sensing, aiming to correlate local physical environments with the specialized biological functions of subcellular compartments. Notably, FNDs have been applied to temperature [24] and radical measurements in mitochondria[16], as well as to probing membrane-associated dynamics and fluidity [22]. These studies highlight the emergence of FND-based ODMR as a powerful platform for elucidating how organelle-specific physicochemical parameters are linked to cellular function.

Despite this progress, quantum sensing within the cell nucleus has remained unexplored. The cell nucleus is a central organelle where physicochemical forces organize chromatin and other nuclear structures to regulate gene expression, highlighting the need to directly elucidate the intranuclear microenvironment through nanoscale measurements of physical parameters. Yet, the application of FNDs for quantum sensing specifically within the nuclear compartment has been limited, because of the fundamental difficulty in simultaneously achieving nuclear accessibility and sufficient ODMR performance.

Cytoplasmic delivery of FNDs is relatively straightforward. For example, FNDs supplemented in the culture medium are readily internalized into cells through endocytosis [25] [26]. In contrast, entry into the nucleus is much more restricted. The nuclear pore complex allows passive diffusion only for particles smaller than ∼30 nm, and indeed nuclear localization has been observed for FNDs below this size [27]. However, such small FNDs produce only weak fluorescence and poor ODMR contrast, making them unsuitable for quantum sensing. On the other hand, FNDs large enough to yield robust ODMR spectra typically measure ∼100 nm in diameter, but these cannot traverse nuclear pores. As a result, ODMR-based quantum sensing within the nucleus has not yet been achieved. Overcoming this barrier requires two key advances: improving ODMR contrast and establishing an effective method for nuclear delivery.

Here, we first optimized the electron-irradiation conditions for sub-100-nm-sized FNDs. Electron irradiation is an established method for creating vacancies and thereby increasing the number of NV centers in nanodiamonds after subsequent annealing [28]. Importantly, NV centers exist in multiple charge states, including the ODMR-active negative NV (NV□, S = 1) and the ODMR-inactive neutral state (NV^0^, S = 1/2) [29]. The relative NV□/NV^0^ population therefore critically influences ODMR signal quality and quantitative sensing performance. Prior studies on electron irradiation of nanodiamonds have primarily optimized NV creation in terms of fluorescence brightness and the NV□/NV^0^ ratio [30]. However, systematic studies focused on optimizing ODMR performance for sensing application using FND remain limited. In this study, we systematically varied the electron-irradiation dose and evaluated the resulting fluorescence and ODMR characteristics, identifying the irradiation conditions best suited for quantum sensing applications.

We next addressed the challenge of establishing a reliable method for intracellular and intranuclear delivery. Microinjection is a well-established technique for introducing molecules or particles directly into individual cells. For FNDs, microinjection has previously been demonstrated in larger biological systems such as embryos, where relatively wide needles can be used [11, 31-34]. However, few reports exist for mammalian cells which require the use of fine capillaries. In these cases, FNDs tend to aggregate and clog the injection needle, preventing efficient delivery. As a consequence, microinjection of FNDs into small mammalian cells, and in particular into the nuclear compartment, has not been achieved. Here, we show that a surface-coating strategy that enhances colloidal stability enables reliable microinjection of FNDs into mammalian cells, including targeted delivery to both the cytoplasm and the nucleus. We further demonstrate field-of-view intranuclear thermometry in living cells.

## 2. Results and Discussion

**Figure 1** illustrates the overall workflow from raw nanodiamond preparation to microinjection into living cells. Commercial type 1b diamond nanoparticles (∼100 nm) were first dispersed in water and subjected to 5 minutes of ultrasonication. The suspension was then kept for 3 minutes, during which large aggregates and poorly dispersible particles settled to the bottom. The supernatant was collected and dried to yield purified nanodiamonds. Although these nanodiamonds contained substitutional nitrogen impurities, they possessed very few vacancies. To generate optically active NV centers, we introduced vacancies via electron irradiation, followed by thermal annealing to pair nitrogen atoms and vacancies. Subsequent air oxidation removed sp^2^ carbon from the particle surface, yielding FNDs. Surface functionalization was then performed to improve dispersibility and compatibility with microinjection. Acid treatment with H□SO□/HNO□ mixture afforded carboxylated FNDs (FND–COOH). Further reaction with glycidol and succinic anhydride produced hyperbranched polyglycerol (HPG)-carboxylated FNDs (FND-HPGCOOH). Dynamic light scattering (DLS) measurements showed that FND–COOH had a median hydrodynamic diameter of 98.9 ± 29.3 nm with a zeta potential of −23.7 mV, whereas FND-HPGCOOH exhibited a median diameter of 136.9 ± 29.1 nm and a zeta potential of −24.0 mV (**Figure S1**). These HPGCOOH-modified FNDs were ultimately employed for intranuclear microinjection and ODMR measurements.

**Figure 1.**
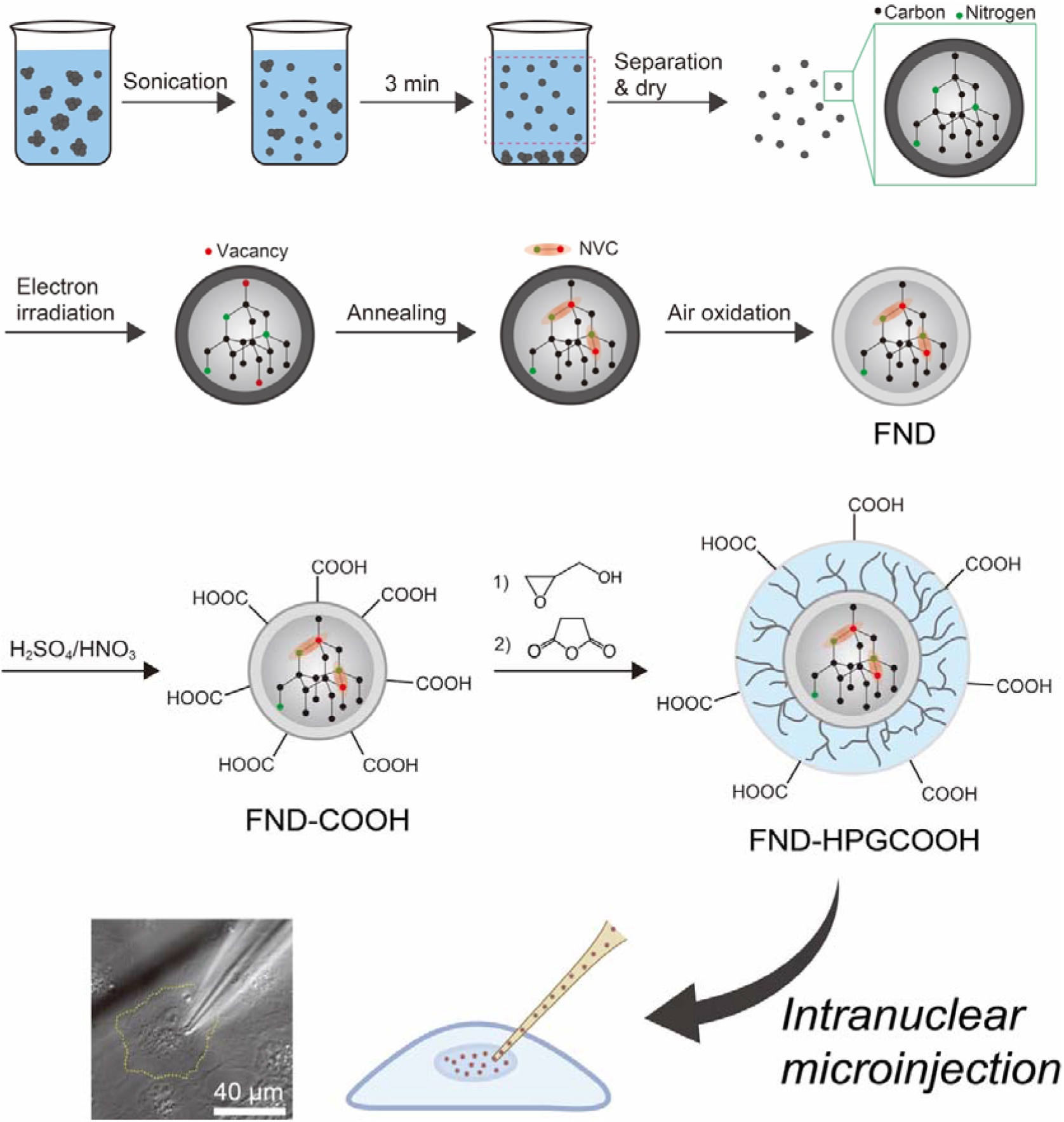
Schematic workflow for preparation and nuclear microinjection of FNDs.

Within this workflow, two steps were particularly critical for successful quantum sensing: optimization of electron irradiation to maximize ODMR contrast, and tailored surface chemistry to suppress aggregation and enable efficient microinjection.

We first evaluated how electron irradiation affects 100-nm FNDs. **Figure 2a** shows the fluorescence spectra of FNDs irradiated at doses of 2, 4, 6, 10, 15, 20, and 30 × 10^1^□ cm□^2^. The peak fluorescence intensity, plotted against dose in **Figure 2b**, increased with irradiation and reached a maximum around 15 × 10^18^ cm□^2^ and then gradually decreased. **Figure 2c** shows the number ratio of NV□ to NV^0^, which was quantified by fluorescence spectral decomposition analysis following the established protocol reported by Alsid et al [35]. This method allows reliable separation of NV□ and NV^0^ contributions and provides a quantitative measure of the NV□/NV^0^ ratio. The ratio was highest at 2 × 10^1^□ cm□^2^ and then gradually decreased with increasing irradiation dose, indicating a progressive shift toward NV^0^-dominant charge states at higher doses. This trend is explained by the availability of excess electrons. NV□ charge stabilization depends on donors from substitutional nitrogen (P1 center). At higher irradiation doses, more vacancies pair with nitrogen atoms to form NV centers, depleting the pool of isolated nitrogen donors that can supply electrons. As a result, the relative abundance of NV□ decreases, while NV^0^ becomes more prominent.

**Figure 2.**
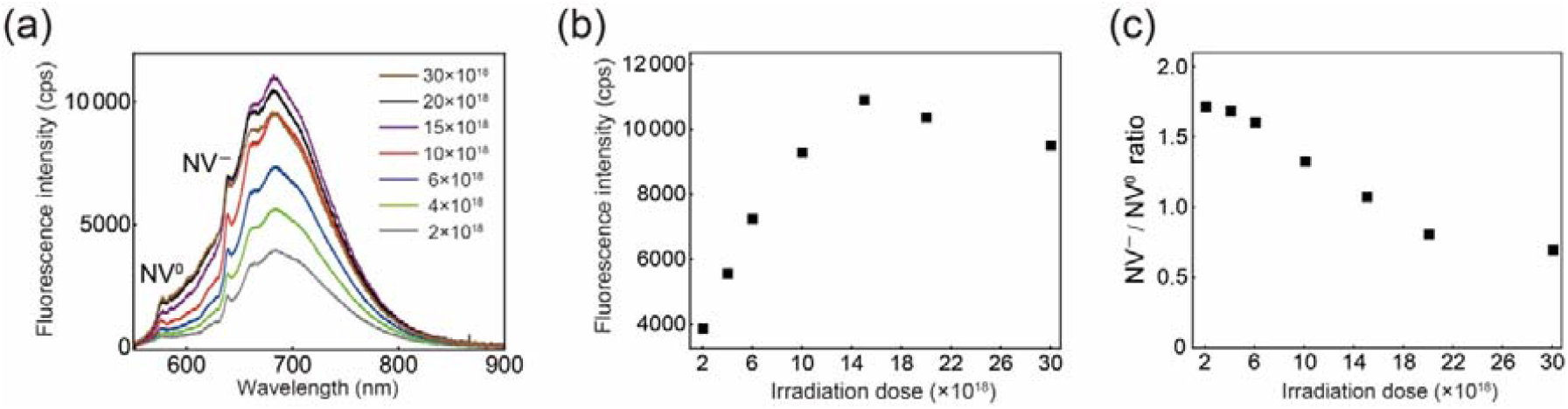
Electron-irradiation dose dependence of fluorescence in FNDs. (a) Representative photoluminescence spectra recorded at various doses. (b) peak fluorescence intensity and (c) number ratio of NV□/NV^0^ as a function of irradiation dose.

Because ODMR is strongly influenced by the total number of NV centers and relative populations of NV□ and NV^0^, we next carried out a detailed analysis of ODMR under the same electron-irradiation conditions. ODMR spectra were obtained from single FND particles and fitted with a sum of two Lorentzian functions (1):

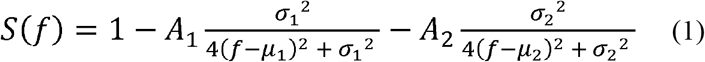

where *A*_1_, *A*_2_ are the signal contrasts, *μ*_1_, *μ*_2_ are the resonance frequencies, *σ*_1_, *σ*_2_ are the linewidth as shown in **Figure 3a**.

**Figure 3.**
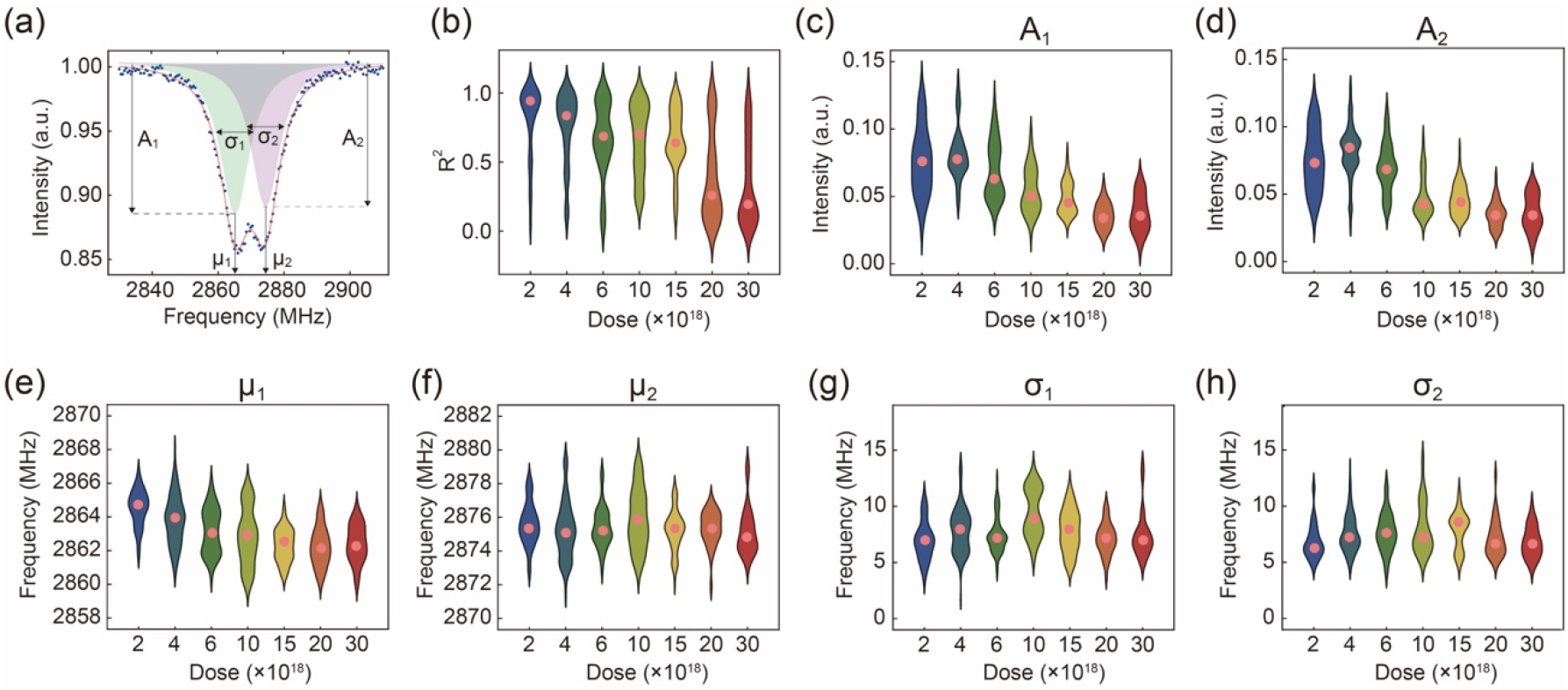
Electron-irradiation dose dependence of ODMR characteristics in single FNDs. (a) Representative ODMR spectrum fitted with a sum of two Lorentzian functions. (b) Coefficient of determination (R^2^) of the Lorentzian fits as a function of irradiation dose. (c, d) Relative contrasts (*A*□, *A*□), (e, f) resonance frequencies (*μ*□, *μ*□), and (g, h) linewidths (*σ*□, *σ*□) extracted from the fits under different irradiation conditions.

The coefficient of determination (R^2^) for the Lorentzian fits decreased progressively with increasing irradiation dose, indicating deteriorating spectral quality (**Figure 3b**). ODMR signal contrast (*A*) increased between 2 × 10^1^□ and 4 × 10^1^□ cm□^2^ but declined at higher doses (**Figure 3c and 3d**). These trends are consistent with the spectral decomposition experiments, where NV□ fraction peaked around lower doses but was suppressed at higher doses (**Figure 2c**). Regarding resonance frequencies (*μ*), *μ*□ exhibited a progressive downshift with increasing dose up to 10 × 10^18^ cm□^2^, beyond which it remained nearly constant (**Figure 3e**). In contrast, *μ*□ was invariant across all doses (**Figure 3f**). The origin of this asymmetric behavior between the two branches is not yet understood. Linewidths (*σ*) showed no systematic dependence on irradiation dose (**Figure 3g and 3h**). Taken together, we identified 4 × 10^1^□ cm□^2^ as the optimal irradiation dose, providing sufficient fluorescence for imaging and the highest ODMR contrast. This dose was adopted in all subsequent experiments.

We next evaluated how surface chemistry influences microinjection performance. Carboxylated FNDs (FND-COOH) show improved dispersibility due to electrostatic repulsion; however, in salt-containing solutions they readily aggregate, and in practice they frequently stuck at the tip of the injection capillary, preventing release (**Figure 4a**). To overcome this, we grafted hyperbranched polyglycerol with terminal carboxyl groups onto the FND surface (FND-HPGCOOH). The hydrophilic HPG shell provides strong steric stabilization, while the additional carboxylates enhance electrostatic repulsion, thereby preventing particle adhesion to the needle tip and enabling smooth release. As anticipated, FND-HPGCOOH dispersed efficiently and could be reproducibly injected without stacking (**Figure 4b, Supplementary movie 1**). We also experimentally screened several dispersibility-enhancing coatings that have been reported for FNDs, including protein, silica, and lipid coatings, by assessing their injectability through fine microinjection capillaries (**Figure S2**).

**Figure 4.**
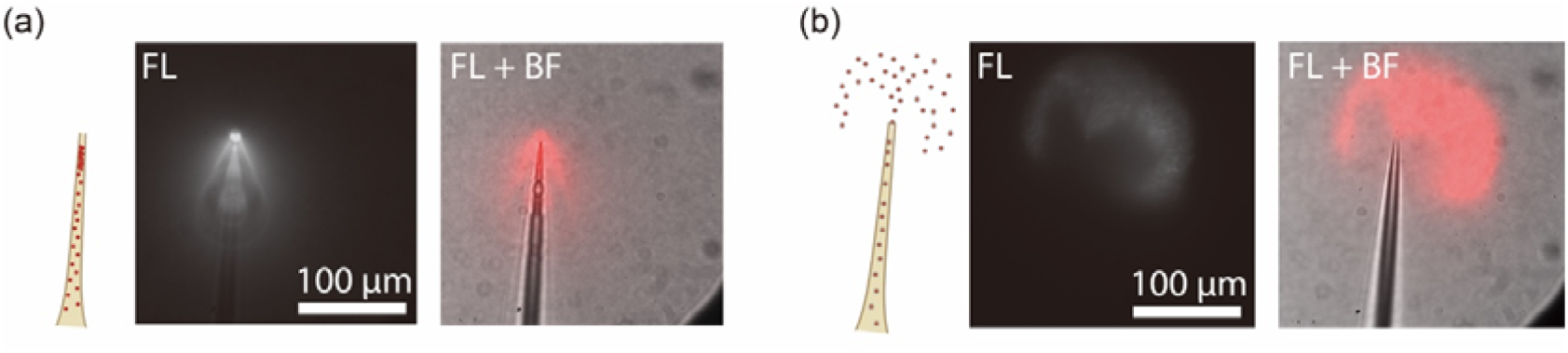
Release behavior of FNDs from injection capillaries. (a) FND-COOH aggregated and stacked at the capillary tip, preventing release. (b) FND-HPGCOOH exhibited stable dispersion and smooth release from the capillary tip, as confirmed by fluorescence (FL) and its merge with bright-field (BF) imaging.

We next performed selective microinjection of FND-HPGCOOH into both the cytoplasm and the nucleus of COS7 cells. The injection site was controlled by adjusting the depth and position of the capillary under microscopic observation, allowing targeted delivery of FNDs to distinct cellular compartments. Representative images show FNDs localized either in the cytoplasm (**Figure 5a and 5b**) or in the nucleus (**Figure 5c and 5d**). Following injection, ODMR spectra were successfully obtained from cytoplasm (**Figure 5e**) and nucleus (**Figure 5f**). Notably, intranuclear FNDs exhibited ODMR spectra with an average contrast of ∼3.5% and signal strength comparable to those measured in the cytoplasm, demonstrating that FNDs can function as sensors even within the nuclear environment. Meanwhile, the signal-to-noise ratio tended to be lower in the nucleus, likely because the smaller nuclear volume contains fewer FNDs within the measurement region, reducing the total number of contributing particles.

**Figure 5.**
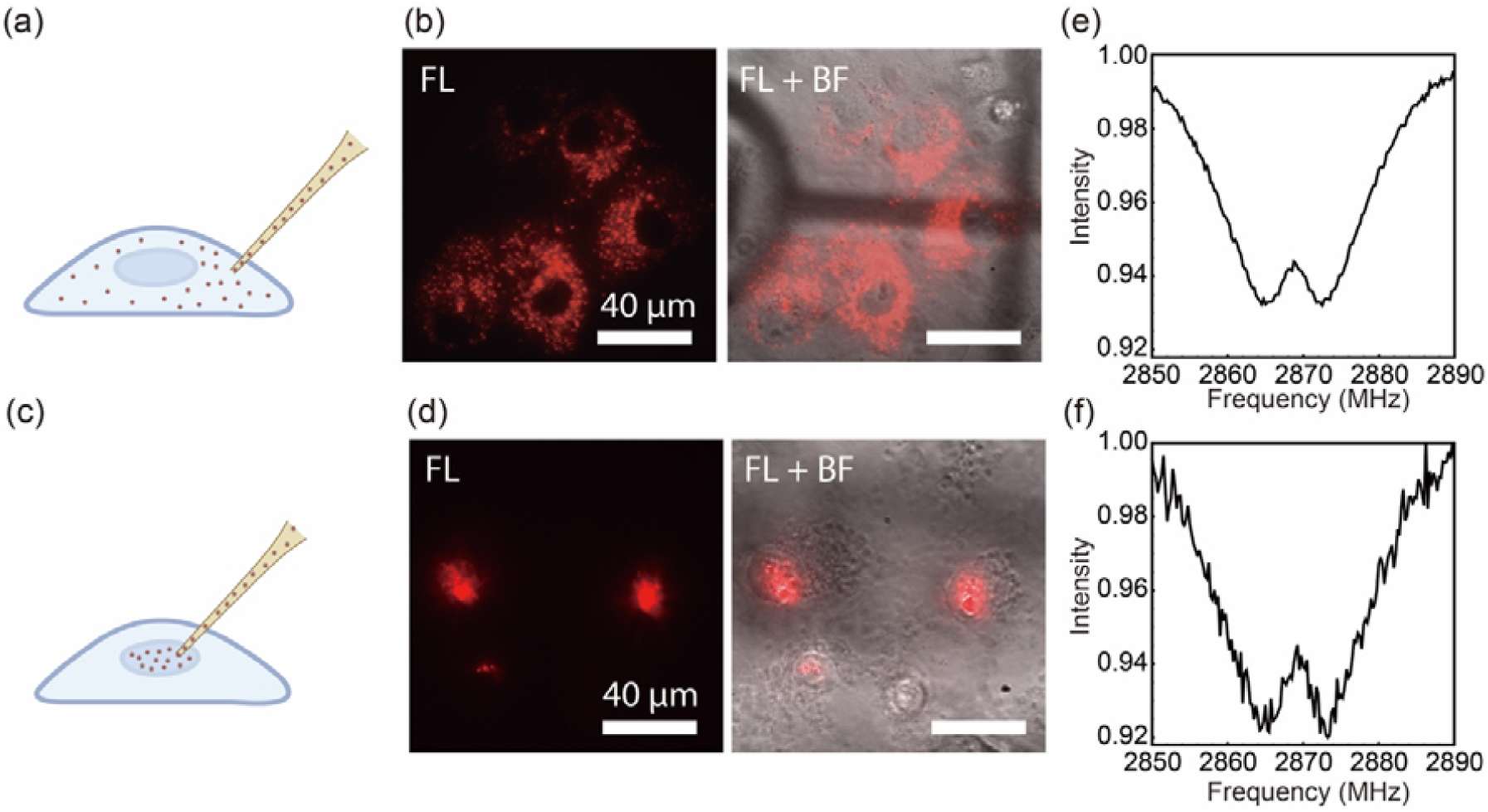
Selective microinjection of FND-HPGCOOH into cytoplasm and nucleus of living COS7 cells. (a,b) Representative confocal fluorescence images showing cytoplasmic localization of injected FNDs. (c,d) Representative confocal fluorescence images showing nuclear localization of injected FNDs. (e, f) ODMR spectra obtained from FNDs in the cytoplasm (e) and nucleus (f).

Finally, we demonstrated field-of-view thermometry in both the cytoplasm and the nucleus using the microinjected FNDs. **Figure 6a** shows a representative fluorescence image containing cells injected either into the cytoplasm or the nucleus. For each region of interest (ROI), we extracted ODMR spectra and determined the center frequency; the corresponding uncertainty was estimated from the covariance matrix of the Lorentzian fitting. We further quantified the frequency shifts induced by controlled changes in the culture-medium temperature, and observed clear, monotonic shifts of the center frequency with temperature for both cytoplasmic (**Figure 6b**) and intranuclear (**Figure 6c**) measurements. The temperature coefficients of the zero-field splitting (ΔD/ΔT) were calculated as −48 kHz °C□^1^ for cytoplasm and −52 kHz °C□^1^ for nuclei. The uncertainties of temperature measurement correspond to ∼1 °C in the cytoplasm and 1–2 °C in the nucleus. At present, the spatial resolution is limited by ensemble averaging, and future integration with targeted or smaller FNDs may further refine intranuclear heterogeneity. Taken together, these measurements attain a new benchmark in intracellular sensing, providing spatial control and quantitative rigor in mapping the thermal environment of the multiple nuclei.

**Figure 6.**
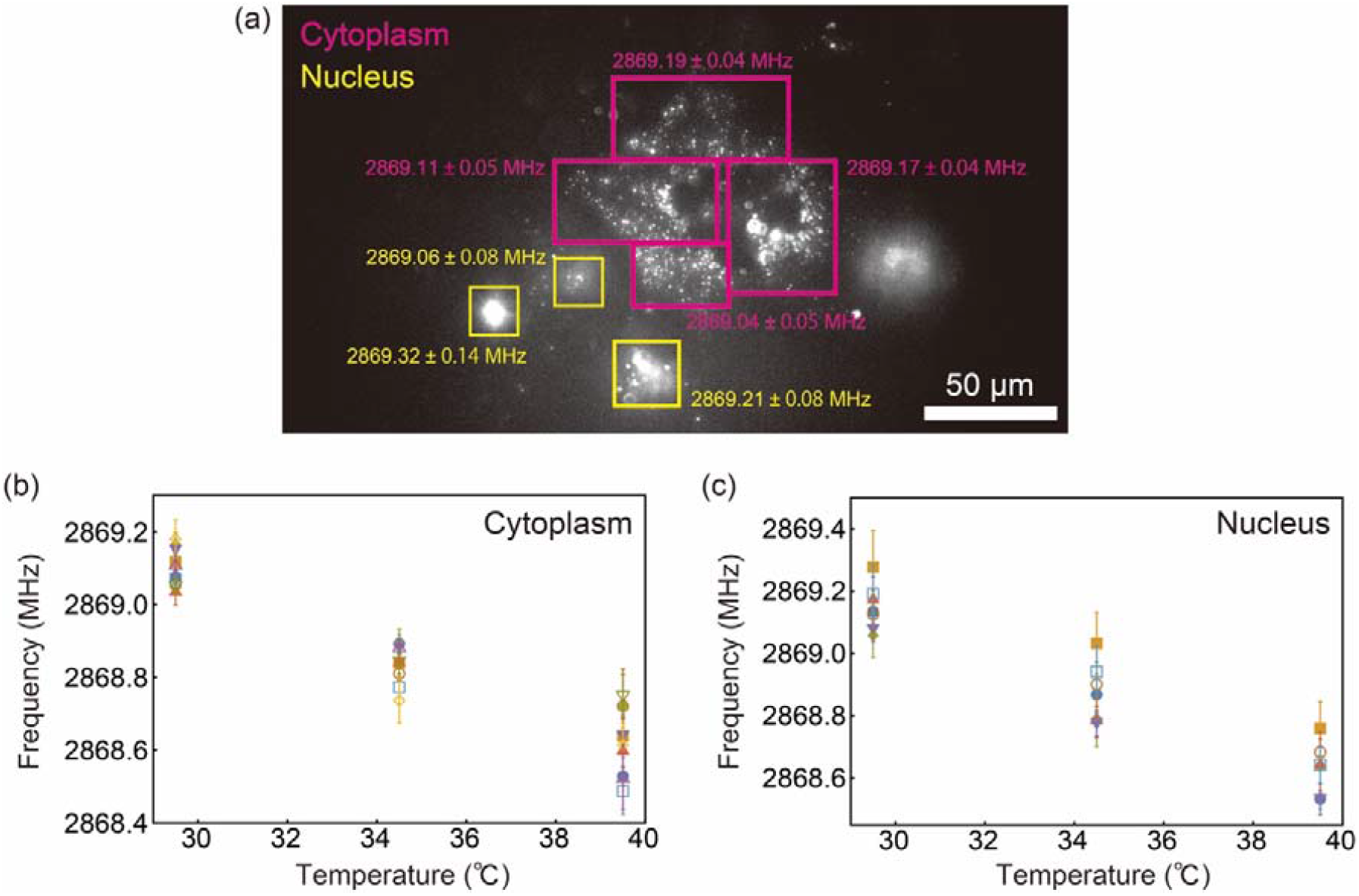
Field-of-view temperature sensing in the cytoplasm and nucleus of living cells using microinjected FNDs. (a) Representative wide-field fluorescence image showing COS7 cells microinjected with FND-HPGCOOH into the cytoplasm (magenta) or the nucleus (yellow). Boxes indicate regions of interest (ROI) used for ODMR analysis, with the extracted center frequencies labeled for each ROI. Temperature dependence of the ODMR center frequency measured from (b) cytoplasmic FNDs and (c) intranuclear FNDs. Error bars represent uncertainties derived from the covariance matrix of Lorentzian

## 3. Conclusion

In this study, we established a robust route to nuclear quantum sensing with fluorescent nanodiamonds (FNDs) by addressing two key bottlenecks: achieving sufficiently strong ODMR performance and ensuring reliable delivery through fine microinjection capillaries. By optimizing the electron-irradiation dose, we identified an irradiation window that balances NV formation with charge-state stability, thereby maximizing ODMR contrast while maintaining fluorescence suitable for microscopy. In parallel, we developed HPGCOOH surface functionalization that suppresses aggregation and prevents clogging or stacking at the capillary tip, enabling smooth and reproducible microinjection. As a proof of concept, we successfully demonstrated field-of-view temperature sensing across multiple cell nuclei simultaneously, providing quantitative and spatially resolved thermal measurement within the genomic environment. Notably, the intranuclear temperature coefficient (ΔD/ΔT) was comparable to that measured in the cytoplasm, confirming that FND-based thermometry remains quantitative in the nucleus.

This intranuclear sensing capability is particularly significant given that the cell nucleus is the center for gene expression and chromatin organization, where local thermal and chemical fluctuations remain poorly understood. Our platform enables the direct interrogation of the chromatin-associated microenvironment, offering a new lens through which to observe the energetic costs of transcription and the thermodynamic regulation of genomic architecture. Furthermore, the compatibility of HPGCOOH-modified FNDs with precise microinjection paves the way for multiparametric quantum metrology, including the simultaneous mapping of temperature, radical species, and local rheology, to dissect the complex mechanochemical signaling pathways that govern cellular identity and disease progression.

## 4. Experimental Section

### Materials

Experimental Details. References are superscripted and appear after the punctuation.

Nanodiamonds (Micron + MDA, 0−100 nm) were obtained from Element Six. Glycidol and tetraethyl orthosilicate (TEOS) were acquired from Sigma-Aldrich. Succinic anhydride, ammonia solution, and tetrahydrofuran (THF) were obtained from Wako Pure Chemical. Bovine serum albumin (BSA) was purchased from Thermo Fisher Scientific. Polyvinylpyrrolidone (PVP) and ethanol were purchased from Nacalai Tesque. 23:2 Diyne PC [DC(8:9)PC] was acquired from Avanti Polar Lipids.

### Preparation of FND

Nanodiamonds were irradiated with a 2 MeV electron beam at doses of 2, 4, 6, 10, 15, 20, and 30 × 10^18^ cm^−2^. The irradiated nanodiamonds were then annealed at 800 °C for 2 h under vacuum. Subsequently, air oxidation was performed at 550 °C for 2 h to obtain FNDs.

### Characterization

The FND solution was excited with a green laser (Compass-215M-50, Coherent, Pennsylvania, USA), and the fluorescence spectra were recorded using a spectrometer (HR4000, Ocean Photonics, Tokyo, Japan). The particle size was measured using a Zetasizer Nano ZSP (Malvern) and surface charge was measured using an ELSZ-1000ZS (Otsuka Electronics, Osaka, Japan).

### Surface modification

FNDs were added to a mixture of sulfuric acid and nitric acid (9:1, v/v) and treated at 70 °C for 3 days. The resulting sample was then reacted in 0.1 M NaOH at 90 °C for 2 h, followed by treatment in 0.1 M HCl at 90 °C for 2 h to produce carboxylated FNDs (FND-COOH). 15 mg of FND-COOH was dispersed in glycidol and washed three times by centrifugation. 4 mL of glycidol was then added to the final pellet, and the mixture was sonicated and stirred at 140 °C for 1 h under an argon atmosphere to synthesize hyperbranched polyglycerol-modified FNDs (FND-HPG). After the reaction, 290 mg of succinic anhydride was added, and the reaction was continued for an additional 1 h. The obtained gel was dissolved in a mixed solution of Milli-Q water and methanol (1:1, v/v) and washed by centrifugation for 30 min. Finally, the product was washed three more times to remove unbound polyglycerol, yielding carboxylated hyperbranched polyglycerol-modified FNDs (FND-HPGCOOH) [36]. 1 mg of FND was dispersed in 10 mL of Milli-Q water and sonicated for 30 min. Subsequently, 1 mg of BSA was added, and the pH of the solution was adjusted to 4.5 using HCl. The mixture was then vortexed for 1 h. To remove unbound proteins, the sample was washed once with Milli-Q water and five times with PBS by centrifugation, yielding FND-protein conjugates [37]. 12 mg of FND and 100 mg of PVP were dispersed in 200 mL of Milli-Q water and sonicated for 3 min. The mixture was stirred for 24 h, followed by three washing cycles with Milli-Q water and one with ethanol by centrifugation. The collected pellet was mixed with 120 *μ*L of TEOS and 500 *μ*L of aqueous ammonia, and ethanol was added to adjust the total volume to 12 mL. After sonication for 1 min, the solution was stirred for 20 h. Subsequently, the product was washed three times with Milli-Q water by centrifugation, yielding silica-coated FNDs (FND-silica) [38]. 1 mg of FND was dispersed in 10 mL of Milli-Q water, and 2 mg of Diyne PC dissolved in 1 mL of THF and 85 *μ*g of cholesterol dissolved in 1 mL of THF were added. The mixture was sonicated for 10 min, and THF was subsequently removed using an evaporator to allow lipid adsorption onto the FND surface. The sample was then irradiated with 254 nm UV light for 90 min to polymerize the lipids. After the reaction, the product was washed three times by centrifugation and filtered through a 0.22 *μ*m syringe filter, yielding lipid-coated FNDs (FND-PCL) [39].

### Cell culture

COS7 cells (JCRB Cell Bank, JCRB9127) were seeded on 60-mm tissue culture dishes (IWAKI) and were cultured in Dulbecco’s Modified Eagle Medium (DMEM, Gibco) supplemented with 10% FBS, penicillin (100 IU/mL), streptomycin (100 *μ*g/mL), L-glutamine (2 mM), sodium pyruvate (1 mM), and MEM at 37°C under a humidified atmosphere containing 5 % CO_2_.

### Dispersion of FND single particles individually over glass using spin coating

The FND suspension was thoroughly sonicated. A mixture was then prepared by combining 15 *μ*L of FNDs (50 *μ*g mL□^1^) with 15 *μ*L of 0.3% (w/w) poly(vinyl alcohol) (PVA). A 30 *μ*L aliquot of the mixture was dispensed onto a 35-mm dish (Matsunami, Japan) rendered hydrophilic by plasma treatment. The dish was mounted in a benchtop centrifuge and spin-coated for 20 s to achieve uniform deposition.

### Microinjections into cells

FND-HPGCOOH (1 mg/mL) was sonicated for 5 min, filtered using a syringe filter with pores of 0.22 *μ*m (Millipore, Burlington, MA, USA), and loaded into a glass capillary needle with an inner tip diameter of 0.5 ± 0.2 *μ*m (Femtotips, Eppendorf, Hamburg, Germany). Pressure was applied with a pneumatic microinjector (FemtoJet 4i, Eppendorf, Hamburg, Germany), and the needle was controlled using a micromanipulator (InjectMan 4, Eppendorf, Hamburg, Germany) to selectively microinject the material into either the cytoplasm or the nucleus.

### ODMR measurements

ODMR spectra were acquired using a custom-built microscope system optimized for quantum sensing. Briefly, a continuous neodymium-doped yttrium aluminum garnet (Nd:YAG) laser at 532 nm illuminated FNDs to initialize and read out the spin state of NV centers on an inverted microscope system (Nikon, Ti-E). Fluorescence from the NV centers was captured by an oil immersion objective lens (Nikon SR HP Apo TIRF, 100×, NA1.49 for FND characterization, Nikon Apo TIRF, 60×, NA 1.49 for the cell experiments), passed through a long-pass filter to cut off the excitation light, and imaged by a camera (EMCCD; Andor, iXon Ultra 888 for FND characterization, qCMOS camera; Hamamatsu, ORCA-Quest 2). A two-turn copper coil with a diameter of approximately 1 mm was placed just above the coverslip to irradiate the sample with MWs at frequencies near the electron spin resonance in NV centers. The MW generated by a MW generator (Agilent, E8257D) was amplified using linear microwave power amplifiers (Mini-Circuits, PAN35-5A and ZHL-16W-43+) and transmitted to the microwave coil via a coaxial cable. For FND characterization, EMCCD exposure time was 20 ms/frame, and total acquisition time was 10 min/measurement. For the cell experiments, qCMOS exposure time was 25 ms/frame, and total acquisition time was 6 min/measurement. The acquisition of camera frames, as well as the timing of microwave and laser pulses, was synchronized via TTL signals generated by a pulse generator (Model 565, Berkeley Nucleonics Corporation).

### Analysis

For FND calibration, the Python Picasso library was used to extract bright spots and determine their fluorescence intensities. In cell experiments, the fluorescence intensity of FNDs within cells was determined by enclosing them within a region of interest (ROI). The ODMR spectra were obtained from the fluorescence intensity ratio during MW ON/OFF states. All fitting procedures were performed using the Python function scipy.optimize.curve_fit. For FND calibration, the ODMR spectra measured at 2830 MHz and 2910 MHz were fitted with a double Lorentzian function to determine the resonance frequencies, contrasts, and full widths at half maximum (FWHM). In cell experiments, the ODMR spectra measured at 2866 MHz and 2972 MHz were fitted with a single Lorentzian function to determine the resonance frequencies. The uncertainty of the fitted parameters was estimated from the diagonal elements of the covariance matrix.

## Acknowledgements

We acknowledge Drs. Ryuji Igarashi, Kiichi Kaminaga, Madoka Suzuki, Kohji Maeda, Yumi Yoshida, Kohki Okabe, and Mr. Hirotaka Okita for their support.

## Supporting Information

Supporting Information is available from the Wiley Online Library or from the author.

## Data Availability Statement

The data that support the findings of this study are available from the corresponding author upon reasonable request.

Received: ((will be filled in by the editorial staff))

Revised: ((will be filled in by the editorial staff))

Published online: ((will be filled in by the editorial staff))

