## Supplementary Information for "Intranuclear Quantum Sensing with Fluorescent Nanodiamond Enabled by Electron-Irradiation and Surface-Chemical Optimization for Microinjection"

Y.S.K.
Department of Biological Sciences, School of Science, The University of Osaka, Machikaneyamacho, Toyonaka, Osaka 560-0043, Japan
Y.S.
Department of Biological Sciences, Graduate School of Science, The University of Osaka, Machikaneyamacho, Toyonaka, Osaka 560-0043, Japan
K.S., S.S., D.F.
Faculty of Molecular Chemistry and Engineering, Kyoto Institute of Technology, Matsugasaki, Sakyo-ku, Kyoto 606-8585, Japan
Y.S., A.G.
Institute for Protein Research, The University of Osaka, Suita, Osaka 565-0871, Japan
H.A., T.O.
Quantum Materials and Application Research Center (QUARC), National Institutes for Quantum Science and Technology (QST), 1233 Watanuki, Takasaki, Gunma 370-1292, Japan
T.O.
Department of Materials Science, Tohoku University, Aoba, Sendai, Miyagi 980-8579, Japan
K.F., Y.H.
Premium Research Institute for Human Metaverse Medicine (WPI-PRIMe), The University of Osaka, Yamadaoka, Suita, Osaka 565-0871, Japan
Y.H.
Center for Quantum Information and Quantum Biology, The University of Osaka, Machikaneyamacho, Toyonaka, Osaka 560-0043, Japan

*
**

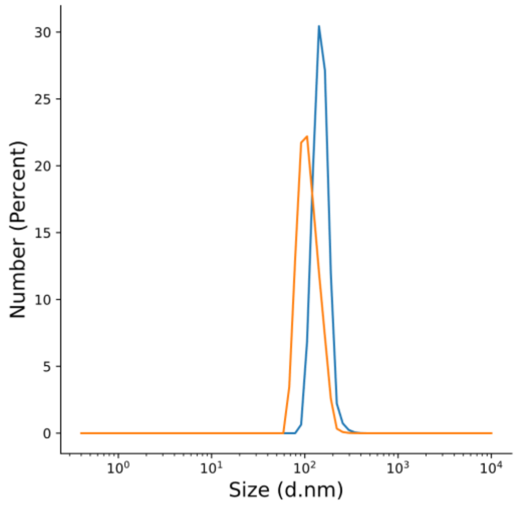


**Figure S1**. Size distributions of FND-COOH (orange) and FND-HPGCOOH (blue) measured by DLS.

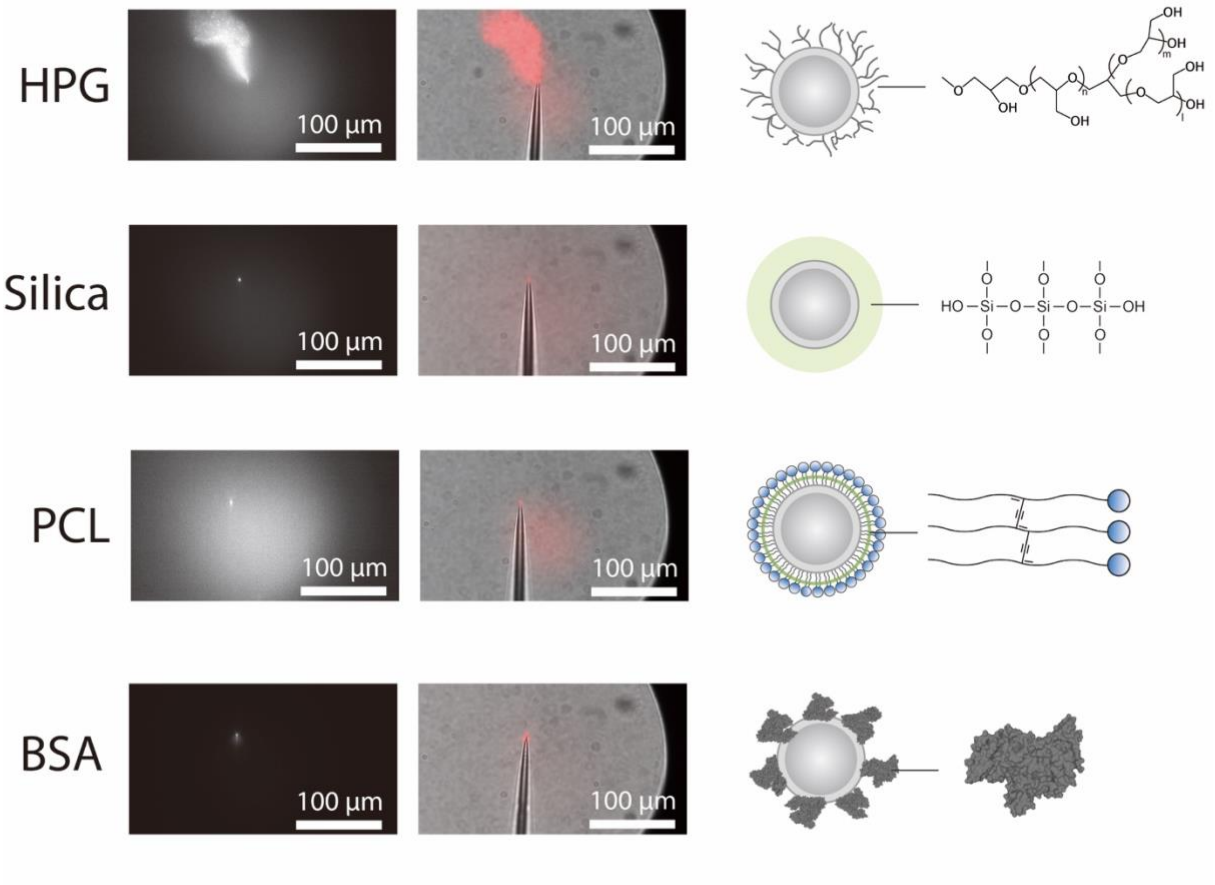


**Figure S2**. Comparison of microinjection compatibility among various FND surface modifications. Fluorescence (left) and merged fluorescence/bright-field (right) images showing the release behavior of FNDs from fine glass capillaries. HPG-modified FNDs exhibited exceptional colloidal stability and were smoothly released from the tip without stacking similar to FND-HPGCOOH. In contrast, FNDs coated with Silica, photo-crosslinked lipid (PCL), or bovine serum albumin (BSA) tended to aggregate and adhere to the inner wall of the needle, leading to clogging. All the coating information is found in elsewhere [S1].
